# IntravChip: a vascularized and perfused microfluidic model of the primary tumor microenvironment to collect intravasated tumor cells

**DOI:** 10.64898/2026.02.19.706805

**Authors:** Marie Floryan, Alessandro Cordiale, Heather Jensen, Julie Chen, Zixian Guo, Vinayak Vinayak, Sina Kheiri, Ritu Raman, Vivek Shenoy, Elena Cambria, Roger Kamm

**Affiliations:** Department of Mechanical Engineering, Massachusetts Institute of Technology; Department of Electronics, Information and Bioengineering, Politecnico di Milano; Department of Biological Engineering, Massachusetts Institute of Technology; Department of Mechanical Engineering and Applied Mechanics, University of Pennsylvania; Department of Materials Science and Engineering, University of Pennsylvania

**Keywords:** intravasation, microfluidics, metastasis, vascular, *in vitro* model, flow

## Abstract

Hematogenous metastasis is initiated when tumor cells (TCs) intravasate into the vasculature, yet intravasation remains poorly understood because it is difficult to observe *in vivo* and intravasated TCs are challenging to isolate. To address these challenges, we developed IntravChip, a continuously perfused microfluidic platform containing a vascularized primary tumor microenvironment (TME) enabling the observation of TC intravasation, and a downstream chamber to collect intravasated TCs. The IntravChip can support a high TC concentration in the TME while maintaining complete vascular perfusion, which we found was necessary to collect intravasated cells. Using MDA-MB-231 breast TCs, we identified an optimal initial TC seeding density that, by day 9, yields a densely populated TME and 100-440 collected intravasated TCs. We validated the IntravChip across several TC types, showing that MDA-MB-231 and MV3 TCs have the highest intravasation rates while MCF-7 TCs have low intravasation efficiency. We also show that the IntravChip is compatible with super-resolution nano-imaging. Our devices enabled high-quality STORM imaging, which revealed that H3K9me3 nanodomains are significantly differentially distributed in intravasated MDA-MB-231 tumor cells compared to those residing in the TME. Finally, the IntravChip was validated as a platform to test the effects of anti-cancer drugs on tumor cells and on the vasculature. We showed that a 5 μM concentration of sorafenib reduced intravasation events by 69% without impacting the morphology of the microvascular networks (MVNs), while a 10 μM concentration led to a significant decrease in vessel diameter. This platform enables quantitative analysis of TC intravasation, collection of intravasated TCs for characterization, and screening of anti-metastatic therapies.

## Introduction

The metastatic cascade consists of a series of stages that tumor cells (TCs) must overcome to spread to distant organs. An early step in hematogenous metastasis, following detachment from the primary tumor, is intravasation, the process by which TCs enter the blood stream. Despite its central role in metastatic dissemination, the dynamics and regulation of intravasation remain poorly understood, largely because these events are transient and rare *in vivo*. *In vivo* observations have identified several mechanisms that promote intravasation and increase metastatic burden, including macrophage-assisted transmigration^1,2^, angiogenesis^3^, vascular remodeling and increased permeability^4^, and increased vessel diameter^5^. Furthermore, *in vivo* and *in vitro* studies have implicated a variety of signaling pathways and molecules in facilitating intravasation, including matrix metalloproteinases (MMPs), epithelial-to-mesenchymal transition (EMT) programs, vascular endothelial growth factor (VEGF), epidermal growth factor receptor (EGFR), and integrin signaling^6^. However, direct observation of intravasation *in vivo* remains technically challenging, relying either on quantifying the number of circulating TCs as a surrogate for intravasation rates^6^ or prolonged high-resolution intravital imaging and three-dimensional reconstruction of densely packed tumor tissue to distinguish intravasated from extravascular cells^2,7,8^. These challenges have motivated the development of more sophisticated *in vitro* models to dissect the mechanisms governing TC intravasation.

While 2D cell culture is widely used to study cancer biology, it does not recapitulate the 3D architecture and function of the primary tumor and the dynamic nature of the metastatic process. To fill this gap, a variety of 3D platforms have recently emerged. Vascularized tumor spheroid models, among other types of models of the tumor microenvironment (TME)^9^, have been used to model the primary TME to study drug transport^10,11^, immune cell recruitment^12^, or immunotherapy efficacy^13^. Endothelial-lined tube models of the primary TME are also prominent and have been used to show that restricting glucose availability decreases TC invasion^14^. Moreover, a wide range of models exist to study the extravasation and later stages of metastasis ^15^, but few models exist to study intravasation. Three types of 3D *in vitro* models have been used to study intravasation: the first type features TCs encapsulated in a hydrogel with one surface of the hydrogel lined with an endothelial monolayer^16–18^; and the second makes use of an endothelial-lined tube embedded in a TC-laden hydrogel^19,20^. In endothelial monolayer-based models, it was shown that TC co-culture with macrophages^18^ or fibroblasts^16^ increase the number of intravasated TCs compared to TC monocultures. Furthermore, disruption of the laminin-rich basement membrane by treatment with methylglyoxal increases the number of extruded TCs into the endothelial monolayer^21^. In this same study, it was also shown that flow along the endothelial monolayer promoted TC-endothelial interactions^21^. While these existing *in vivo* and *in vitro* approaches have provided valuable insight into the cellular and molecular contributors to intravasation, there remains a critical need for experimental platforms that enable direct observation of intravasation, quantitative measurements of intravasation rates, and recovery of intravasated cells for further analysis.

To address this gap, we developed the IntravChip, a microfluidic platform that incorporates a perfused primary TME and a downstream chamber to collect intravasated TCs for analysis. We characterize the platform by testing the effect of flow, TC density, and TC type on TC intravasation, and further demonstrate its compatibility with super-resolution imaging to assess chromatin structure within TCs upon intravasation. Finally, we show how an anti-cancer drug affects the structure of the vascularized TME and the ability of TCs to intravasate. This vascularized, perfused microfluidic primary tumor model supports mechanistic dissection of intravasation and direct access to intravasated cells, thus enhancing our understanding of metastasis and providing a powerful tool for preclinical anti-cancer drug screening.

## Methods

### Cell culture

Immortalized human umbilical vein endothelial cells (ECs) (Lonza, CC: 2935, immortalized and transfected to express blue fluorescent protein as previously described^22^), and human primary normal lung fibroblasts (FBs) (Lifeline, FC-0049) were used in this study. ECs were cultured in VascuLife VEGF Endothelial Medium Complete Kit with one quarter of the supplied Heparin Sulfate LifeFactor (VascuLife, Lifeline, LL-0003) and FBs were cultured in FibroLife S2 Fibroblast Medium Complete Kit (Lifeline, LL-0011). MDA-MB-231 (CRM-HTB-26, ATCC) transfected to express cytoplasmic tdTomato and tagged with H2BC11-EGFP, as previously described^23^, SN12-PM6 expressing red fluorescent protein (RFP), MCF7 expressing mCherry, and MV3 expressing cytoplasmic tdTomato and tagged with H2BC11-EGFP were cultured in Dulbecco’s Modified Eagle’s Medium (ThermoFischer) with 10% fetal bovine serum and 100 U mL−1 penicillin and 0.1 mg mL−1 streptomycin (Sigma-Aldrich).

### Microfluidic device and pump fabrication

The IntravChip is composed of a cylindrical gel region, which houses the primary TME, flanked by two media channels on either side, and ports to enable connection to a microfluidic pump to provide continuous, recirculating unidirectional flow (**Fig. 1 A, B**). Microfluidic devices and pumps were assembled as previously described^24,25^. In brief, polydimethylsiloxane (PDMS, Dow Corning Sylgard 184, Ellsworth Adhesives) mixed at a 10:1 ratio of base to cross-linker was, degassed and, cured overnight in a mold at 65°C. The PDMS was cut into individual devices or pumps and ports were punched using biopsy punches (Integra Miltex). Devices and pumps were sterilized in an autoclave. Glass coverslips (VWR, #1) were disinfected in 100% EtOH and dried. Silicone membranes (LMS, Amazon) were cut to size and biopsy punches (Integra Miltex) were used to punch ports. Membranes were then sterilized in an autoclave. The devices and #1 coverslips were exposed to plasma (Herrick Plasma), bonded together, and placed in a 75°C oven overnight. The pumps were assembled, first the bottom of the pump was bonded to the silicone membrane, next the half of the pump was bonded to the bottom, finally, the system was placed in a 75°C oven overnight, as previously described.

**Figure 1.**
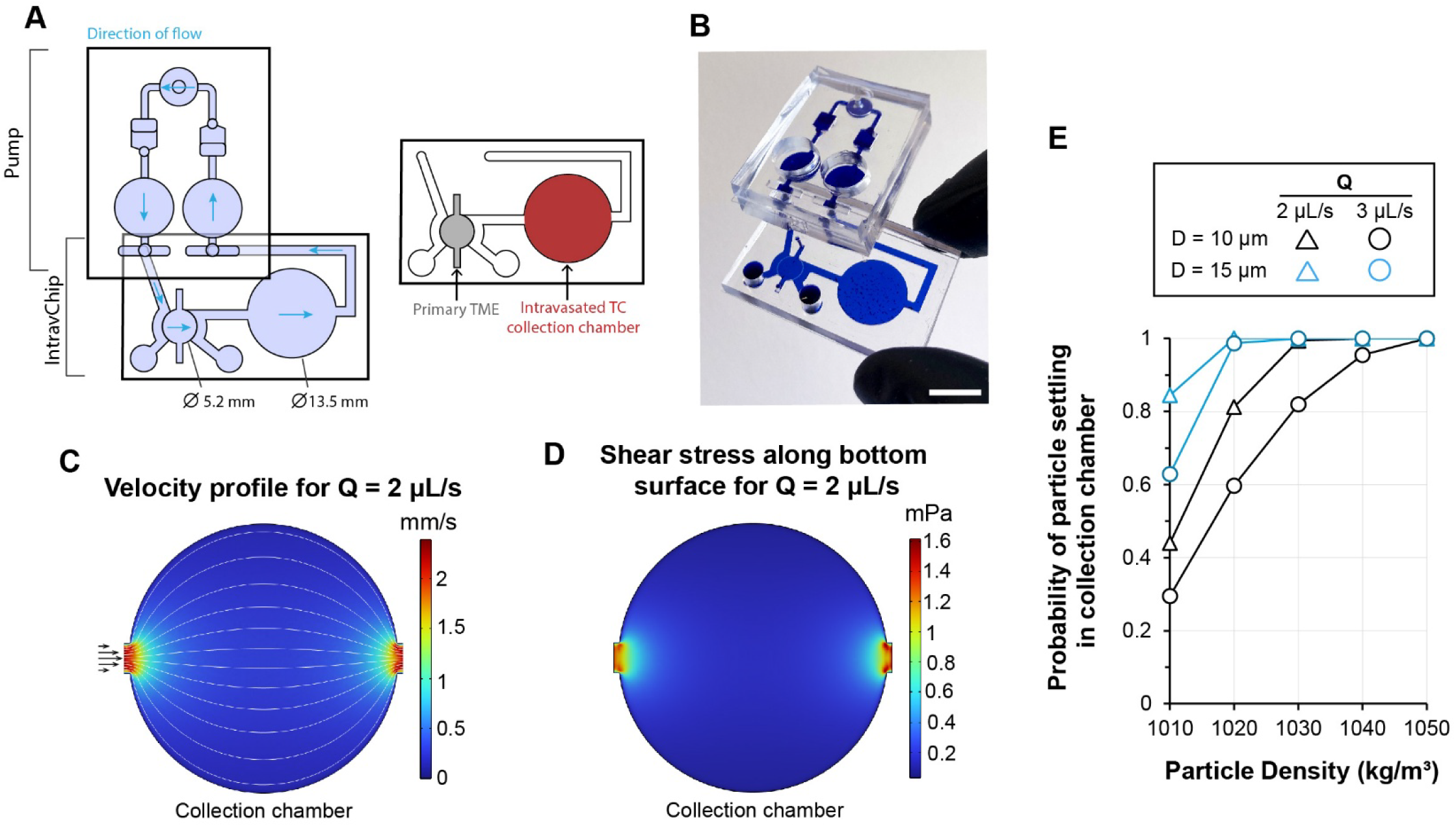
Description of the IntravChip and computational simulation results used in designing the IntravChip. (A) Schematic of the IntravChip connected to a microfluidic pump. Blue arrows indicate the direction of fluid flow and light purple indicates perfused media channels and tissue regions. The grey region is the primary TME and the red region is the intravasated TC collection chamber. The chip height is 500 μm. (B) Picture of the IntravChip connected to the microfluidic pump. Scale bar is 1 cm. (C) Velocity field and streamlines along the center (z = 250 μm) of the collection chamber for a bulk flow rate of 2 μL/s. (D) Wall shear stresses along the bottom surface of the collection chamber for a bulk flow rate of 2 μL/s. (E) Particle tracing simulations were used to identify probabilities of collecting particles (representing TCs) of varying densities, diameters, and bulk flow rates.

### Microfluidic device seeding and culture

ECs at passage 10 and FBs at passage 5 were used for all experiments. ECs and FBs were detached using Accutase (Sigma), pelleted, and resuspended in cold Vasculife supplemented with 4 U mL^-1^ thrombin (Sigma). ECs were resuspended at a concentration of 26×10^6^ cells mL^-1^ and FBs were resuspended at a concentration of 4×10^6^ cells mL^-1^, concentrations selected to provide a large vascular surface area and thereby maximizing intravasation rates. The resuspended ECs and FBs were then mixed together at a 1:1 ratio and placed on ice. TCs were detached with Accutase (Millipore Sigma) and resuspended at a concentration of 357,000 cells-mL^-1^ in cold PBS to yield an initial seeding number of 1000 TCs. Lower initial TC seeding numbers were attained by using either a 10x or 100x dilution with cold PBS. The TC suspension was then mixed with 5 mg/mL freshly thawed fibrinogen (Sigma) at a 3:5 volume ratio of fibrinogen:TC suspension and placed on ice. The EC-FB mixture was mixed with the TC-fibrinogen mixture at a 1:1 ratio and 15 μL were injected into the gel channel of the IntravChip. This process was repeated for all devices. The devices were placed in a humidified incubator for 12 minutes to allow the hydrogel to polymerize. Warm VascuLife was then added to the media channels, and devices were replaced in the incubator. Media changes were performed daily. For static devices media was changed daily. For devices with flow, a microfluidic pump was connected to devices on day 4 and the pneumatic input was set to 5 kPa, which corresponds to aa flow rate of 2 μL/s for a MVN with a hydraulic resistance of 1e11 Pa-s-m^-3^ by previous calibration of the pump system^24^. The medium was changed on days 5 and 7, as previously described^24^. This flow rate was used as a boundary condition in a previously described computational model^26^ to estimate the vessel fluid velocities.

### Imaging

The primary TME was imaged on a confocal microscope (Olympus FV1000) using a 4× objective at a resolution of 640 pixels. The collection chamber was imaged on the same system as 16 ROIs arranged in a 4×4 grid using a 4× objective at 512 pixel resolution. Vessel permeability was measured as previously described^27^. In brief, 0.1 mg mL^-1^ FITC conjugated 70 kDa dextran (Sigma) was added to the media channels, after which the device was quickly transferred to an imaging stage and imaged at times t = 0 min and t = 12 min using a 10X objective at 640 pixel resolution on a confocal microscope (Olympus FV1000).

### Image Analysis

Vessel morphology was assessed as previously described^24^. Briefly, maximum projections of z-stacks produced 2D images which were then binarized using Trainable Weka Segmentation (v4.0.0) in ImageJ (NIH). The projected vessel area was quantified, the image was skeletonized, and the skeleton was analyzed. The average diameter was calculated as 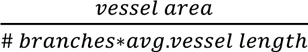. Similarly, TC area in the TME, a surrogate for cell number, was similarly quantified from 2D binarized maximum projection images using Trainable Weka Segmentation in ImageJ. The number of TCs in the collection chamber was determined using ImageJ to stitch z-stack images forming a 4×4 grid. A maximum projection was then applied, the boundary of the collection chamber was marked using the bright field channel, and the image was binarized using the Triangle method, ensuring the background signal was minimal. A watershed was applied and the Analyze Particles plugin was used to count particles larger than 450 μm^2^. Imaris (Oxford Instruments, v.11), was used to generate 3D renderings of z-stacks.

### Computer simulations for vessels and collection chamber

Vessel velocities were determined through simulations using micro-Vascular Evaluation System (muVes) in MATLAB R2025b as previously described^26^. Briefly, a threshold was applied to 2D maximum projected images of fluorescent dextran-perfused vessels which were then input into the 2D muVes environment in MATLAB. Pressure boundary conditions were imposed at the media channels corresponding to a 2 μL/s bulk flow rate and the vessel velocities were computed. Fluid flow through the collection chamber was simulated using COMSOL Multiphysics v.6.3. A Solidworks CAD file of the collection chamber was imported into a 3D COMSOL environment. The laminar fluid interface was employed for steady flow to determine the velocity profile, using parameter values for water at 37°C with a density of 1000.7 kg/m^3^ and a viscosity of 0.791 mPa-s. A constant flow rate of 2 or 3 μL/s was imposed at the inlet. The generated flow field was then used for particle tracking. At time t=0, 1000 particles simulating TCs were released at the entrance to the collection chamber at a spatial distribution proportional to the velocity profile. Gravity force, drag force, and a freeze wall condition were assigned to the particles. Particle diameters of10 and 15 μm and densities of 1010-1050 kg/m^3^ were used^28^ The simulation time was selected to ensure that no particles were moving by the end of the run, at which point the number of particles contacting the bottom surface was counted. The probability of collecting TCs in the collection chamber was calculated using the following formula:

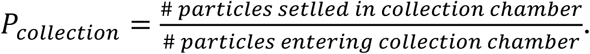

### Immunofluorescence staining

Cells cultured within the microfluidic intravasation device were fixed with 4% paraformaldehyde in PBS for 24 h at 4°C. Following fixation, samples were washed with PBS three times. For UEA-1 lectin staining, samples were incubated for 20 min at room temperature with DyLight 649 UEA-1 lectin (VectorLabs, DL-1068-1, 1:200 dilution in PBS) and subsequently washed three times with PBS. Samples used for STORM were incubated for 1 h at room temperature in permeabilization and blocking buffer consisting of 10% donkey serum, 0.3% Triton X-100, and 0.1% BSA in 1× PBS. Cells were then incubated with a primary antibody against H3K9me3 (Invitrogen, 720093; 1:100 dilution) at 4°C overnight. Devices were then washed with PBS at room temperature for at least 6 h with hourly buffer exchanges. Samples were subsequently incubated with a Alexa647-conjugated secondary antibody (Abcam, ab150075; 1:100 dilution) at 4°C overnight. The following day, samples were washed in PBS for 1 h at room temperature. All buffer exchanges were performed using a syringe or pipette tip connected to the outlet of the microfluidic channel to avoid introducing air bubbles. After washing, devices were kept in PBS until STORM imaging.

### STORM imaging of microfluidic device

STORM imaging was performed using an oxygen-scavenging imaging buffer as previously described^29^. The buffer contained 10 mM cysteamine (MEA; Sigma-Aldrich, 30070-50G), 0.5 mg/mL glucose oxidase, 40 mg/mL catalase, and 10% (w/v) glucose in PBS, enabling efficient photo switching of Alexa647 fluorophores. Prior to imaging, samples were incubated in the imaging buffer for at least 30 min to ensure thorough exchange of PBS and to fully immerse the imaging region. Super-resolution imaging was performed on a commercial Oxford NanoImager (ONI) platform equipped with a ×100, 1.4 NA oil-immersion objective and a Hamamatsu ORCA-Flash sCMOS camera. STORM datasets were acquired with a 15 ms exposure time for 30,000 consecutive frames using 647 nm excitation. Single-molecule localizations were extracted using ONI’s native acquisition and reconstruction software. Resulting localization coordinates were further processed and analyzed using in-house MATLAB scripts as previously described^30–32^. MDA-MB-231 cells were engineered to express H2B–GFP, which permitted selective identification of tumor cells in the presence of unlabeled ECs, ensuring that quantification reflected only tumor cell chromatin architecture. Intravasated TCs were sparsely distributed in the collection chamber, and we conducted extensive searching across the entire imaging area to locate and acquire STORM datasets from all GFP and Alexa 647 positive nuclei.

### Statistical analysis

Statistical analysis and data visualization was conducted using GraphPad Prism 8 (v10.2.0). Normality tests were performed using the Shapiro-Wilk test and the Kolmogorov-Smirnov test. For detection of outliers the ROUT test was used for identifying multiple outliers and the Grubbs’ test was used for identifying one outlier. For assessment of statistical significance, Student’s t-tests, Welch’s t-tests, ordinary one-way ANOVAs, or two-way ANOVAs were performed. Statistical significance was assigned as follows: p>0.05 (ns), p<0.05 (*), p<0.01 (**), p<0.001 (***), p<0.0001 (****).

## Results

### Particle tracking simulations informed the design of the IntravChip collection chamber

The IntravChip was designed to optimize the collection efficiency of intravasated TCs carried by flow from an engineered TME. The intravasated TCs are carried by the flow into the downstream collection chamber where they settled and adhered to the bottom surface (**Fig. 1 A, B**). The minimum distance between the edge of the primary TME and the edge of the collection chamber was 8.3 mm to ensure that TCs would not migrate along the glass into the collection chamber^33–37^ (**Supp. Fig. 1**). By requiring that the settling time for a TC that enters the collection chamber be less than the residence time, we can establish the following requirement for the minimum radius of the collection chamber:

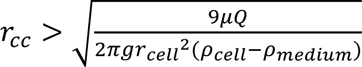, where *ρ*_cell_ is the cell density, *ρ* is the medium density, *μ* is the medium viscosity, *Q* is the flow rate, *r_cell_* is the radius of a cell, and *g* is gravity. **See Supp. Fig. 2** for details. For a cell density of 1.056 g/mL^28,38,39^ and fluid properties for medium at 37°C^28^, a cell diameter of 14.3 μm^38^, and a *Q* of 1.2 μL/s, the minimum radius of the collection chamber was estimated at 6.75 mm.

Computational particle tracking simulations confirmed that using these dimensions, TCs settle to the bottom with flow rates up to 3 μL/s (**Fig. 1 C**). Particle tracking simulations were used to assess the effect of cell density and diameter on collection probability. As the cell density even within a single cell type can vary by about +/-0.005 g/mL^40^, we examined the effect on settling at different densities. For a flow rate of 2 μL/s or 3 μL/s, the simulation predicted that all particles with a diameter of 10-15 μm and a density of 1050 kg/m^3^ would settle to the bottom surface (**Fig. 1 C**). For a 15 μm diameter particle, the settling probability only decreased below 100% for particle densities of below 1020 kg/m^3^, while for a 10 μm diameter particle the settling probability decreased for particle densities below 1030 kg/m^3^ (**Fig. 1 C**). Furthermore, particles in the simulation settle in a radial pattern from the inlet, whereas experimentally the TCs were more dispersed (**Supp. Fig. 3**). This may be due to TC rolling along the coverslip prior to adhesion and TC migration under flow^33–37^. Of the collected TCs, 23% were found in clusters with an average number of ∼5 TCs per cluster (**Supp. Fig. 3**). These results confirm that the collection chamber geometry effectively captured TCs of >10 μm diameter with densities >1040 kg/m^3^ at a bulk flow rate of 2 μL/s. At this flow rate, TCs initially enter the collection chamber with an average velocity of about 2.5 mm/s and then slow to <0.5 mm/s throughout most of the chamber (**Fig. 1 D**). Furthermore, the wall shear stress along the bottom plane is <0.6 mPa over the majority of the collection chamber (**Fig. 1 E**), well below reported limits of cell detachment^41^, suggesting that once the cell settles to the bottom and adheres, it should remain attached to the surface. These results show that the chosen flow rate and collection chamber geometry used in the IntravChip are optimal to collect intravasated TCs from continuously recirculating medium.

### High magnification images of the primary TME show TC-EC interactions

On day 4 after seeding the ECs, FBs, and highly metastatic MDA-MB-231 TCs in a fibrin gel, when the ECs had connected to form a perfusable microvascular network (MVN), the pump was connected and the system was continuously perfused until day 9 (**Fig. 2 A**). Formation of the TME was observed over a 9-day period and was imaged at high magnification on day 9 to investigate the physical relationship between the TCs and vasculature. The TCs significantly proliferated in the TME, increasing from ∼2% area coverage on day 1 up to ∼40% on day 9 (**Fig. 2 B**). Fluorescently tagged dextran (70 kDa) was used to confirm perfusability (**Fig. 2 C**) and assess vessel permeability (**Fig. 2 D**). Although the TC-laden MVNs tended to have higher permeability than the TC-free MVNs, the difference was not statistically significant (**Fig. 2 D**). Furthermore, vessels in TC-laden TMEs tended to have a slightly smaller average vessel diameter and reduced vessel area coverage, however no significant differences in MVN morphology were observed between TC-free and TC-laden TMEs (**Supp. Fig. 4**). The bulk flow rate employed in the IntravChip experiments (2 μL/s) corresponds to an average vessel fluid velocity in the TME of 5.3 mm/s (**Fig. 2 E**), somewhat higher than previously reported vessel velocities for mammary breast tumor microvasculature^42^. The TME tissue was examined in confocal images leading to several general observations. TCs were often observed spreading directly on the basal side of the endothelium and occasionally appearing to lie within the endothelium (**Fig. 2 F**). However, higher resolution imaging techniques such as scanning electron microscopy may be needed to better visualize these cell-cell interactions. TCs were also observed directly lining the vessel lumen (**Fig. 2 G**), resembling mosaic vessels (defined as vessel lumens partially covered by TCs), previously observed in vivo^43,44^. We also observed what appeared to be a TC in the process of intravasation, indicated by the bleb-like protrusion of the TC into the luminal space (**Fig. 2 H**). These are exceptionally rare events that could, however, be further investigated through subsequent studies using the IntravChip. These results demonstrate that the IntravChip houses a perfusable vascularized TME with a tight endothelial barrier and that TC-EC interactions implicated in intravasation can be visualized.

**Figure 2.**
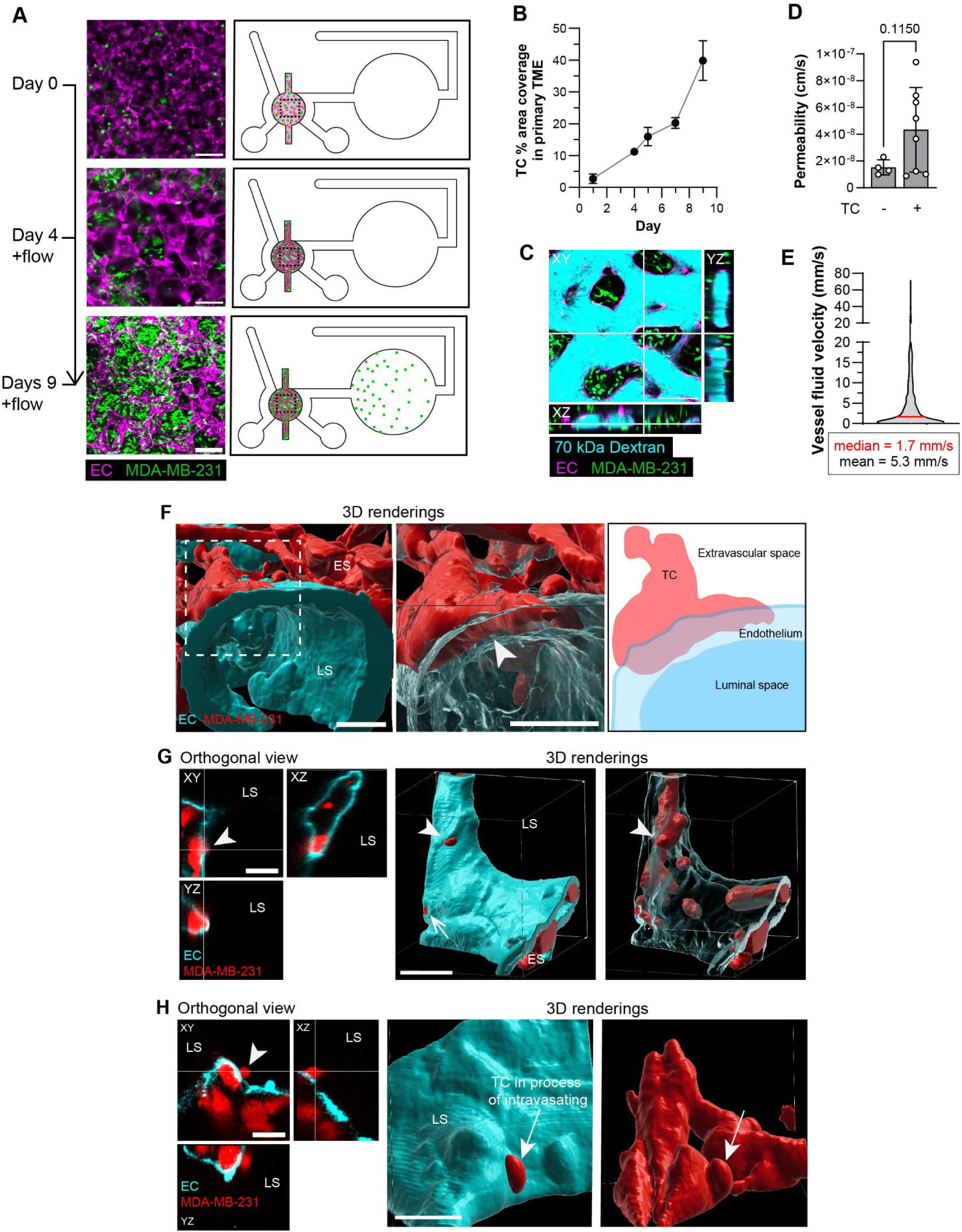
Experimental timeline and characterization of the TME region. (A) Experimental timeline. ECs, FBs, and TCs are seeded in the primary TME region on day 0. Vessels form by day 4 and continuous perfusion begins and continues until day 9. TCs settle in the collection chamber between days 4 and 9. Scale bar is 200 μm. (B) 2D TC % area coverage from day 1 to day 9. n = 5-7 devices for each timepoint. (C) Cross section view of vessels perfused with FITC 70 kDa Dextran (cyan). TCs (green) surround vessels. Scale bar is 200 μm. (D) Vascular permeability measurements for day 7 MVNs seeded with or without TCs. n = 4-8 devices. (E) Truncated violin plot of vessel fluid velocities. n = 1095 vessels from n = 4 devices. Red line indicates the median vessel velocity. (F-H) Orthogonal views and 3D renderings of confocal images using a 60X objective, scale bars are 20 μm. LS: luminal space, ES: extravascular space. (F) MDA-MB-231 cells lying along and within the basal surface of the endothelium (arrowhead). Middle panel is zoomed in on a TC of interest, and the endothelium surface is transparent. Right panel is a 2D graphic of the middle panel. (G) Orthogonal view (left panel) and 3D rendering (middle and right panels) of TC partially lining the endothelium in the luminal space. Arrowhead is pointing at the same TC across views. Arrow corresponds to a second site where a TC is partially lining the endothelium. The middle and right panels are of the same view but with different endothelial surface transparencies. (H) Orthogonal view (left panel) and 3D rendering (middle and right panels) of a TC in the process of intravasating (arrow). The middle and right panels are of the same view; the right panel only shows the TC surface rendering. B, D: graphs show mean +/-sd.

### Continuous flow yields higher vascular coverage, TC proliferation, and numbers of collected TCs

To examine the effect of flow on MDA-MB-231 TC intravasation, we compared continuously perfused systems to ones maintained under static conditions. We have previously shown that continuous perfusion promotes significant remodeling of engineered MVNs over time^24^, but the role of flow on MVNs in an engineered TME has not been described. Samples from both conditions were grown under static culture for the first four days, after which either the pump was connected to the flow samples for continuous perfusion until day 9 or samples remained under static conditions. Noting that the vessels in these MVNs have an elliptical cross section with the major axis in the XY-plane (**Fig. 2 C**, ^24^), vessel diameters measured from projected z-stack images represent maximum vessel diameters. By day 9, the MVNs in the flow condition occupied 15% more area (**Fig. 3 A, B**), had 41% larger average diameters (**Fig. 3 C**), 25% longer vessels (**Fig. 3 D**), and a lower vessel density (**Fig. 3 E**). Note that the average vessel density in the IntravChip of 74 vessels/mm^2^ falls within the reported range of micro vessel density from breast cancer patient biopsies^45^. We also assessed the TC area coverage in the TME as a surrogate for cell number and found that the perfused TME showed 30% greater TC coverage compared to static (**Fig. 3 F**), suggesting higher proliferation rates. We observed a significant increase in the number of collected TCs in the perfused condition compared to static (**Fig. 3 G**), reflecting the fact that flow is the predominant mechanism by which cells can enter the collection chamber. On average 240 intravasated TCs were collected under flow while only 12 were collected in static condition (**Fig. 3 G**), likely a consequence of transient flows generated during media changes. These results demonstrate that flow has a significant effect on vessel morphology, TC proliferation, and is required to collect intravasated TCs.

**Figure 3.**
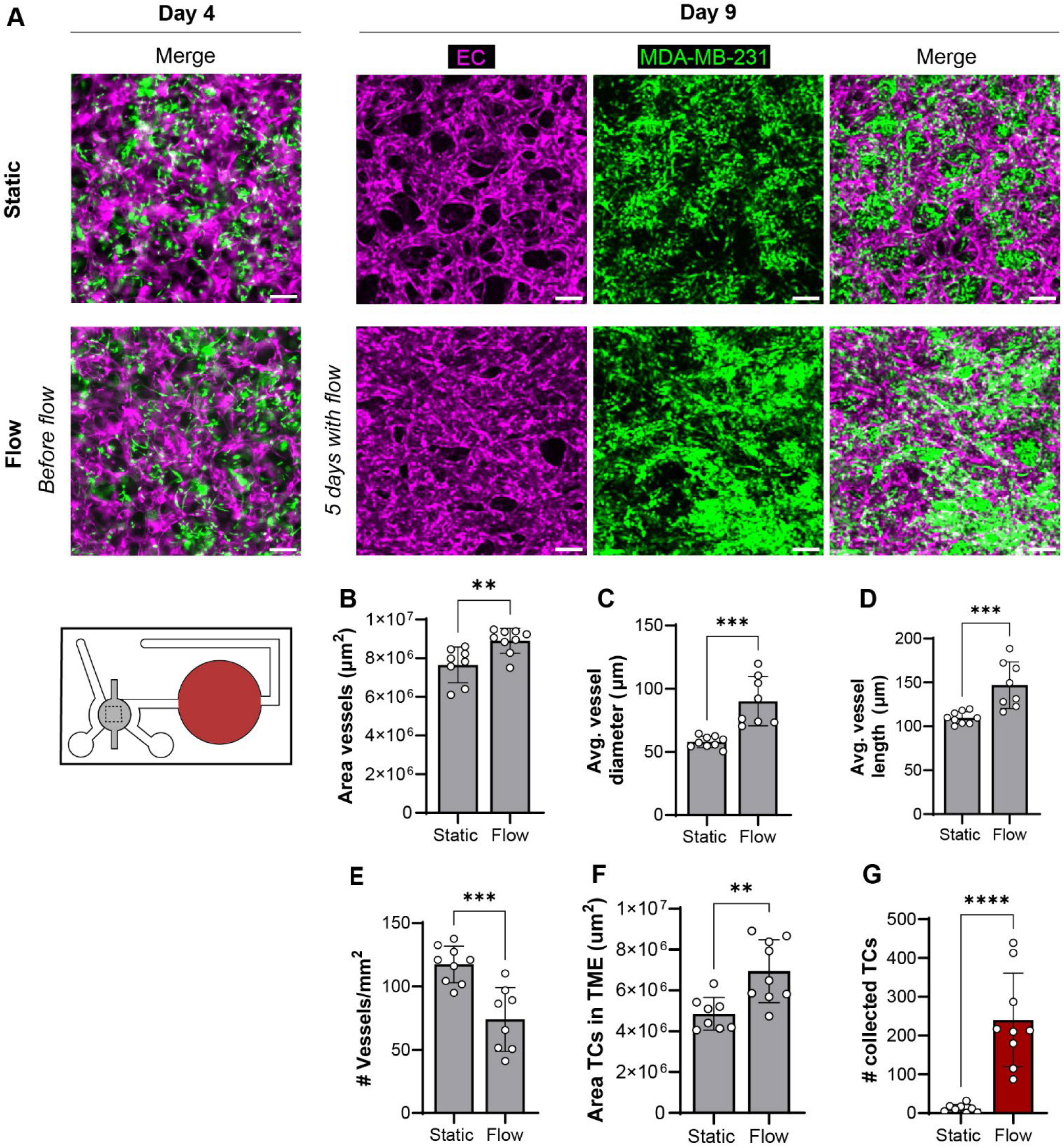
Effect of flow on vascular morphology, primary TME, and number of intravasated cells. (A) Representative maximum projection images of the TME from static and flow conditions on day 4 (before flow) and on day S (static or 5 days with flow). IntravChip diagram colorings. Grey corresponds to the primary TME and red corresponds to the collection chamber. Scale bar is 200 μm. (B-E) MVN morphology quantifications for (B) vessel area coverage, (C) average vessel diameter, (D) average vessel length, and (E) vessel density. (F) TC area coverage, a surrogate for TC number, in the primary TME. (G) Absolute number of collected TCs. n = 8-S devices. Graphs show mean +/-sd. Asterisks indicate p-value ranges as follows: *p < 0.05, **p < 0.01, ***p < 0.001, ****p < 0.0001.

### Intravasation potential depends on primary TME TC density

We next examined the effects of TC density in the primary TME and culture duration on intravasation rates for MDA-MB-231 TCs. Our earlier findings showed that an initial TC density of 1000 TCs/device yields a highly dense TME (**Fig. 2, 3**). Therefore, the other TC densities we investigated were lower, corresponding to 10 and 100 TCs per TME. While the 1000 TC density yields widespread coverage of TCs in the TME, the TME in the lower densities is characterized by localized TC colonies and many regions without TCs (**Fig. 4 A**). The vessel area across all conditions and timepoints remained similar (**Fig. 4 B**). The TC area in the TME increased with increasing initial TC density (**Fig. 4 C**), however TC proliferation rates over time differed across the TC densities. The TC area increased by about 50% in both the 10 TC and 100 TC groups between days 7 and 9 but only increased by about 21% in the 1000 TC group (**Fig. 4 C**). The 1000 TC group had the highest average number of collected cells, 77 on day 7 and 102 on day 9, while fewer than 10 TCs were collected in the 10 TC and 100 TC on days 7 and 9 (**Fig. 4 D**). Furthermore, the number of collected intravasated TCs increased from day 7 to day 9 in the 1000 TC group. Interestingly, the 10 TC group had the highest number of intravasated collected TCs normalized to the TC area in the TME (**Fig. 4 E**), which may indicate that the TCs more efficiently intravasate at very low densities. Based on these results, we chose to analyze all subsequent experiments at day 9 and use initial seeding conditions of 1000 TCs to maximize the absolute number of collected intravasated TCs.

**Figure 4.**
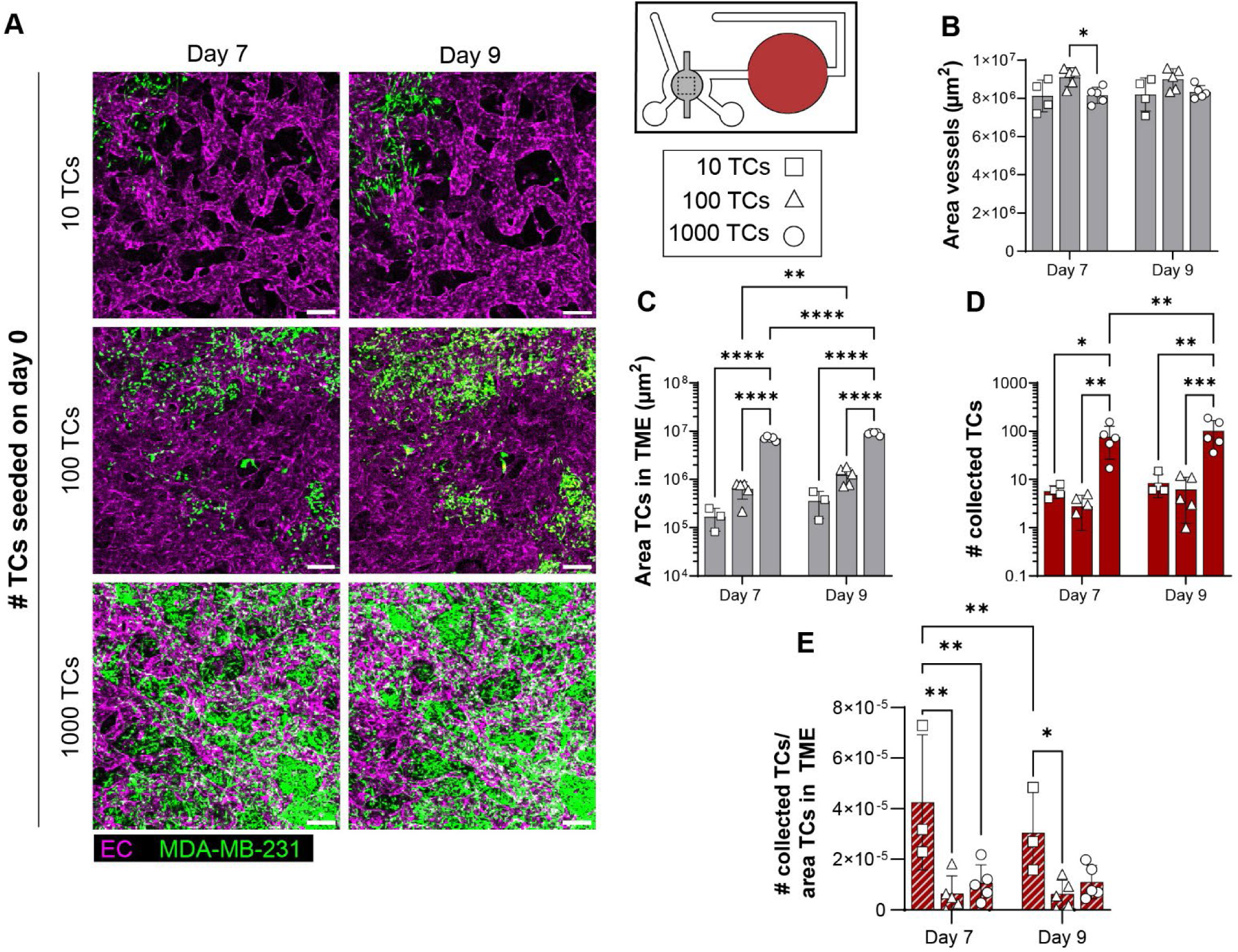
Primary TME TC area coverage and number of collected intravasated TCs as a function of initial TC seeding. (A) Representative maximum projection images of the TME on days 7 and S from TMEs that were initially seeded with either 10, 100, or 1000 TCs per device at day 0. Scale bar is 200 μm. (B-D) Quantification of (B) vessel area coverage, (C) TC area coverage from 2D project z-stacks, (D) number of collected TCs, and (E) number of collected TCs normalized to the area of TCs in the TME. n = 3-5 devices. Graphs show mean +/-sd. Asterisks indicate p-value ranges as follows: *p < 0.05, **p < 0.01, ***p < 0.001, ****p < 0.0001.

### Intravasation potential depends on TC type

We then assessed the model’s capability to distinguish the primary TME structure and intravasation potential across different TC types. We seeded TMEs with an initial TC density of 1000 cells per device using either highly metastatic MDA-MB-231, highly metastatic melanoma line MV3, highly metastatic renal cell carcinoma line SN12-PM6, or low metastatic MCF7 to confirm that the IntravChip can capture expected differences in intravasation rates across TC types. The structure of the primary TME varied across TC types. The MCF7 TME consisted of few, highly compact TC clusters, while the MDA-MB-231 and MV3 were more uniformly dispersed, and the SN12-PM6 exhibited a combination of the two (**Fig. 5 A**). Interestingly, vessel area coverage did not differ across TC types (**Fig. 5 B**), potentially indicating that the vessel remodeling due to continuous flow may be the dominant factor in determining vessel area coverage. By day 9, TC area coverage varied greatly across TC types, with MDA-MB-231 and MV3 TCs showing the largest TC area, followed by the SN12-PM6, and the MCF7 exhibiting the lowest coverage (**Fig 5 C**). A similar trend was observed in the absolute and normalized number of collected TCs (**Fig. 5 D, E**).

**Figure 5.**
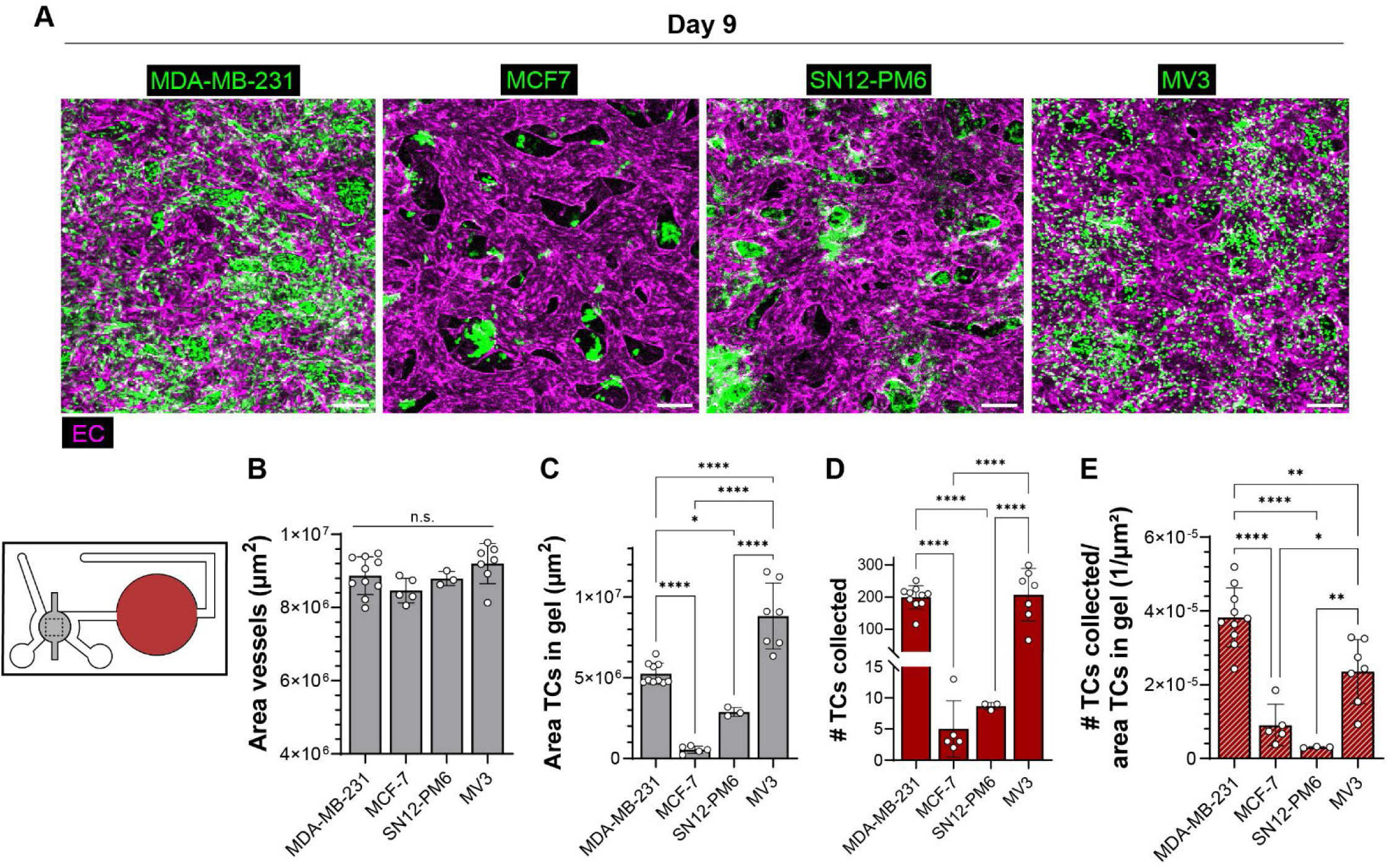
Intravasation potential across different TC types. (A) Representative maximum projection images of the TME with MDA-MB-231, MCF7, SN12-PMC, or MV3 TCs. Scale bar is 200 μm. (B-C) Quantification of (B) 2D projected vessel area coverage, (C) 2D projected TC area coverage in the TME. (D-E) The intravasation rate was assessed by (D) the absolute number of collected TCs and (E) the number of intravasated TCs normalized to the TC area coverage in the TME. n = 3-10 devices. Graphs show mean +/-sd. Asterisks indicate p-value ranges as follows: *p < 0.05, **p < 0.01, ***p < 0.001, ****p < 0.0001, and n.s. is not significant.

Although the SN12-PM6 are known to extravasate with moderate efficiency and comparably to MDA-MB-231 TCs^46^, the results here indicate that their ability to intravasate is comparable to that of the weakly metastatic MCF7. These results confirm that the IntravChip can recapitulate trends in invasiveness across TC types described in literature and can be used to assess intravasation rates. All following studies use only the MDA-MB-231 TCs.

### STORM imaging reveals preserved heterochromatin abundance but dispersed organization after intravasation

To show that the IntravChip can be used to characterize mechanisms at different scales, including the nanoscale, we investigated how intravasation influences nano-scale chromatin organization. Mechanical constraints and microenvironmental cues are known to profoundly reshape nuclear architecture of TCs, including the heterochromatin organization^47,48^. To study heterochromatin organization we performed STochastic Optical Reconstruction Microscopy (STORM) and quantitative analysis on H3K9me3-labeled MDA-MB-231 cells cultured on glass as a control, within the primary TME, and in the collection chamber in the IntravChip.

The super-resolution images showed that TCs cultured directly on glass exhibited significantly higher H3K9me3 localization densities compared to TCs in the IntravChip **(Fig. 6 A, B)**. However, total H3K9me3 localization counts per nucleus did not differ significantly between primary TME and intravasated TCs **(Fig. 6 A, B)**. Despite these differences in abundance, STORM imaging revealed pronounced changes in the nanoscale organization of heterochromatin following TC culture in the primary TME and intravasation **(Fig. 6 C, D)**. To quantify these structural differences, H3K9me3 nanodomains were identified using Density-Based Spatial Clustering of Applications with Noise (DBSCAN) clustering of single-molecule localizations. TCs cultured on glass displayed large domains, with an average radius of 162.7 nm, while TCs in the primary TME TCs contained smaller domains, with an average radius of 126.7 nm, and intravasated TCs contained even smaller domains, with an average radius of 81.7 nm **(Fig. 6 C)**. TCs on glass control and intravasated TCs exhibited comparable numbers of H3K9me3 nanodomains per nucleus, whereas primary TCs in TME contained substantially fewer nanodomains **(Fig. 6 D)**. Together, these results show that the IntravChip can capture nano-scale heterochromatin remodeling upon intravasation.

**Figure 6.**
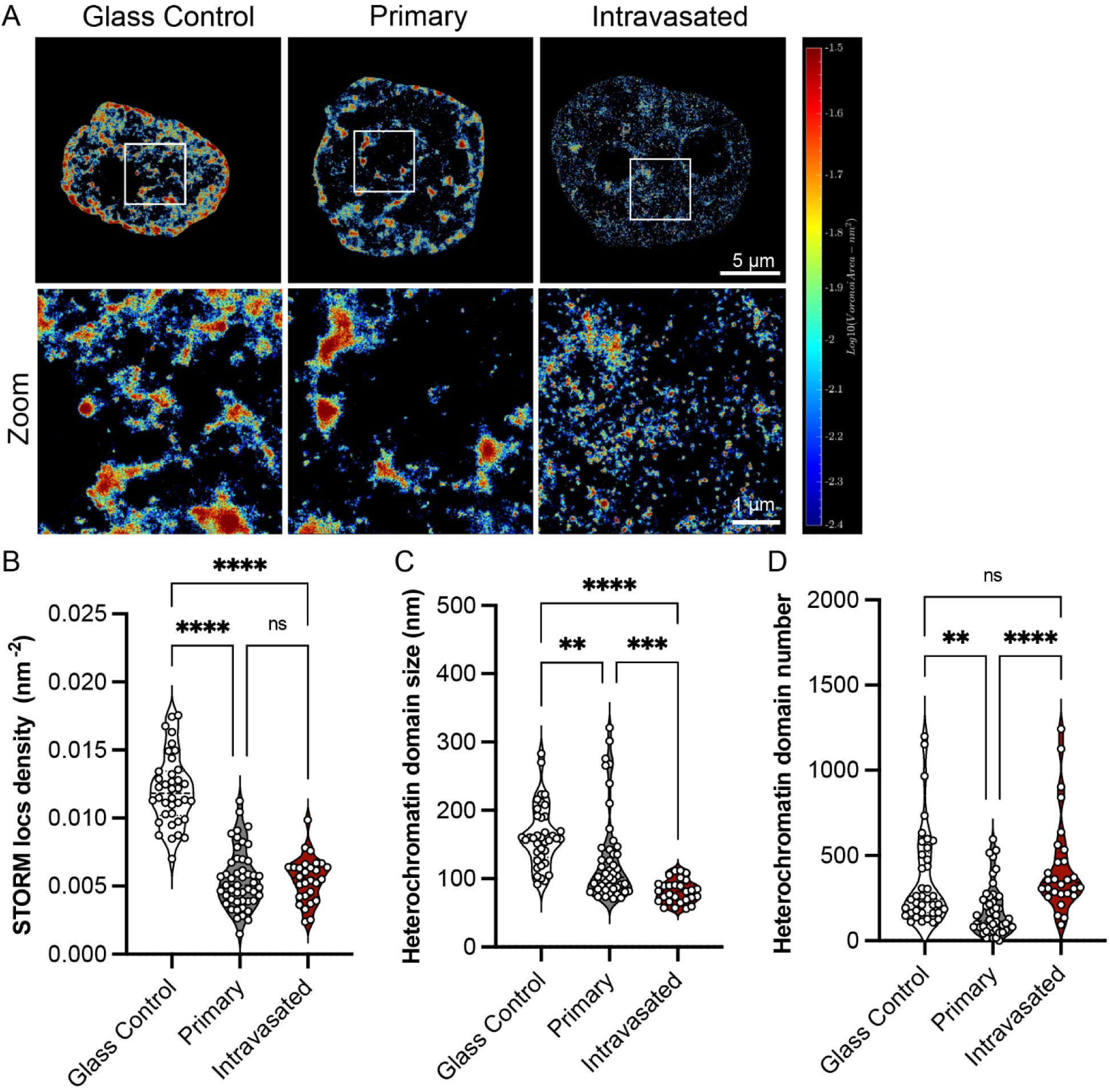
Chromatin nano-imaging in IntravChip. (A) Voronoi density maps of H3K9me3 localizations on glass control TCs, primary TCs and intravasated TCs, reconstructed from STORM imaging. Each polygon represents the local single molecular localization density surrounding an individual localization, with red colors indicating higher density. Scale bar is 5μm. (B) Quantification of H3KSme3 localization density per unit nuclear area for glass control, primary and intravasated TCs. (C) Quantification of heterochromatin nanodomain size derived from clustering analysis of H3K9me3 localizations. (D) Total number of heterochromatic nanodomains per nucleus for glass control, primary TME, and intravasated TCs. All experiments were performed using at least three devices. Sample sizes were n=38, n = 48 and n = 28 TCs for glass control, primary and intravasated TCs, respectively. Statistical significance was assessed using two-sided Student’s t tests. Asterisks indicate p-value ranges as follows: *p < 0.05, **p < 0.01, ***p < 0.001, ****p < 0.0001; n.s., not significant (p ≥ 0.05).

### Sorafenib treatment reduces intravasation events

We treated the model with sorafenib to confirm that is sensitive to anticancer drugs and possibly useful as a drug screening platform. Sorafenib is a an oral multikinase inhibitor with antiproliferative, antiangiogenic, and proapoptotic effects on tumors and is used to slow the growth of several types of refractory solid tumors^49^. It has been shown to reduce intravasation events in an *in vitro* platform^16^, making it a suitable benchmark for platform validation.

The MDA-MB-231 TME was treated with two concentrations of sorafenib, 5 μM and 10 μM, selected based on prior studies that examining sorafenib effects on MDA-MB-231 TCs^50–52^. Sorafenib and vehicle control (0.01% DMSO) treatments began on day 4 with the initiation of flow, followed by media changes and replenishment of sorafenib or vehicle on days 5 and 7 (**Fig. 7 A**). The TME and the collection chamber were then analyzed on day 9 (**Fig. 7 A**). Across all conditions the vessels remained fully connected (**Fig. 7 B**). However, vessels treated with 10 μM sorafenib had reduced vessel area (**Fig. 7 C**) and smaller diameters (**Fig. 7 D**) compared to control, while the morphology of the vessels treated with 5 μM sorafenib was unaffected compared to control. There were no significant differences in average vessel length (**Fig. 7 E**) or vessel density (**Fig. 7 F**) across conditions. TC area coverage in the primary TME decreased with increasing concentration of sorafenib, decreasing by 35% and 59% for 5 μM and 10 μM, respectively (**Fig.7 B, G**), confirming that our model can recapitulate 2D *in vitro*^50,52^ and *in vivo*^53^ observations of sorafenib treatment reducing TC proliferation and tumor size. Furthermore, both concentrations of sorafenib similarly reduced intravasation events by 83% compared to vehicle control (**Fig. 7 H**), and when normalizing against the TC area coverage in the TME, the 5 μM sorafenib treatment reduced intravasation events by 75% (**Fig. 7 I**), indicating that decreased TC proliferation does not alone account for the reduction in collected TCs. Together, these results demonstrate that there is a sorafenib dose range that is sufficient to reduce TC proliferation and intravasation events without causing vessel regression in the TME. They further show that the IntravChip captures drug effects on the TME TCs, TME vessels, and intravasation, supporting its potential use as an assay to assess anti-intravasation drugs.

**Figure 7.**
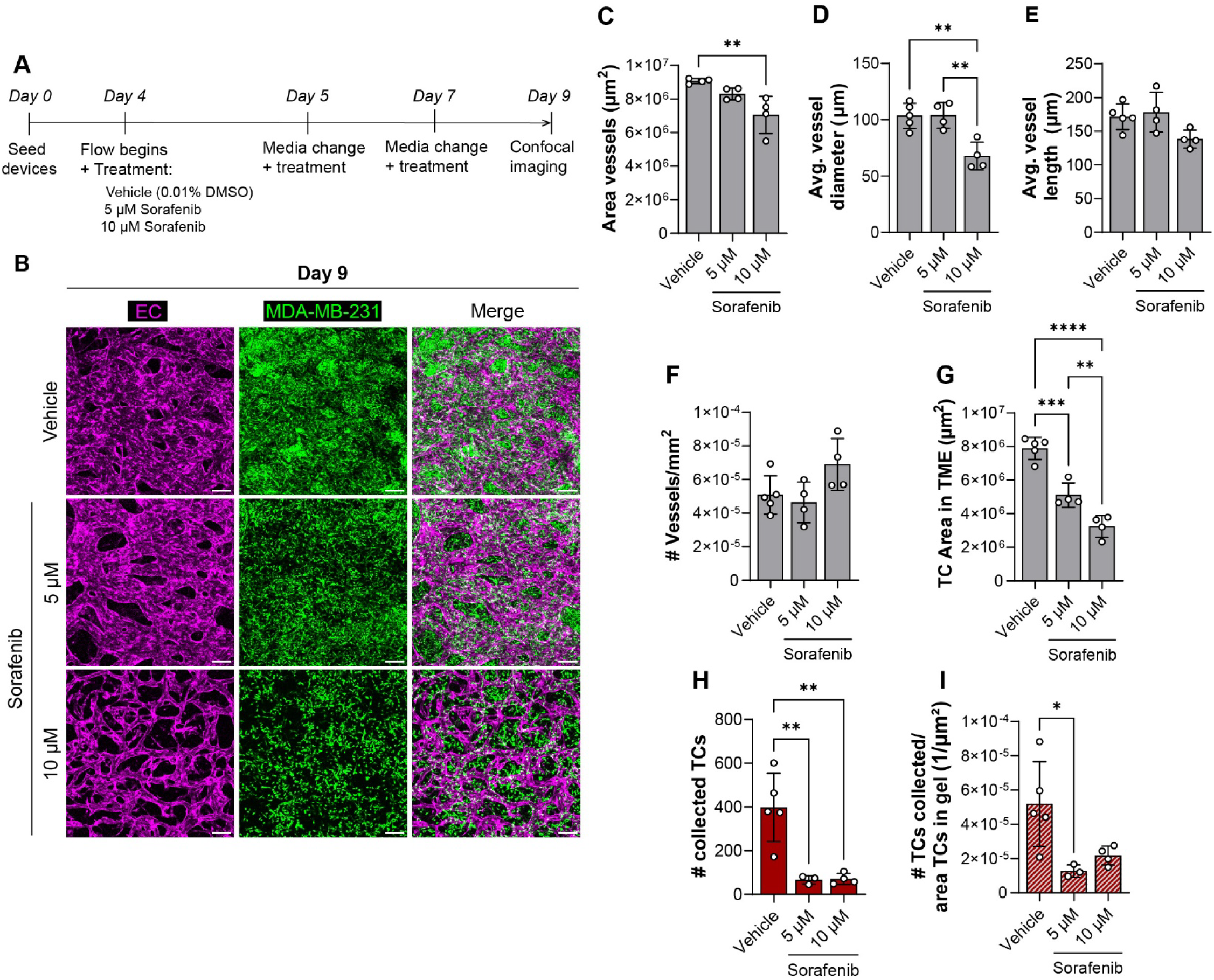
Dose-dependent effect of sorafenib on vasculature, TC area coverage, and intravasation. (A) Overview of the experimental timeline. (B) Representative maximum projection images from day 9 of vehicle, and 5 μM or 10 μM sorafenib-treated MDA-MB-231 TMEs. Scale bar is 200 μm. (C-F) Quantification of MVN morphology parameters: (C) 2D projected vessel area coverage, (D) average vessel diameter, (E) average vessel length, and (F) vessel density. (G) Quantification of the 2D projected TC area coverage in the TME. (H-I) Intravasation rate was assessed by (H) the absolute number of collected TCs and (I) the relative number of collected TCs normalized to the TC area coverage in the TME. n = 4-5 devices. Graphs show mean +/- sd. Asterisks indicate p-value ranges as follows: *p < 0.05, **p < 0.01, ***p < 0.001, ****p < 0.0001.

## Discussion

While there has been progress in the development of preclinical models of the primary TME for studying the early stages of the metastatic cascade^16–18,54^, direct assessment of intravasated TCs or intravasation rates remains a challenge. In this study, we presented a novel system, the IntravChip, consisting of a 3D-vascularized primary TME with continuous perfusion to model the early stages of metastatic dissemination and to collect intravasated TCs for subsequent analysis. We show that the IntravChip can be used to generate vascularized and continuously perfused TMEs with varying TC densities and TC types while using flow to collect intravasated TCs to assess intravasation rates. Using the IntravChip, we show that 1) TC interactions with the endothelial monolayer can be readily visualized, 2) the MVN morphology is more strongly affected by intravascular flow than the presence of TCs, 3) differences in TC invasiveness from TMEs with varying TC densities and TC types can be clearly identified, 4) intravasated TCs have more accessible chromatin domains than TCs at the primary tumor site, and 5) sorafenib reduces the occurrence of intravasation events.

At the site of the primary tumor, the IntravChip provides the opportunity to gain new insights into the details of the intravasation process. Once the TC enters the lumen, it may reside there for a time, break loose and enter the circulation, or attempt to migrate inside the lumen possibly coinciding with the generation of an intravascular matrix. It is also possible that fibroblasts from the TME accompany the TC in this process, although we have no direct evidence of that. Although the scarcity of events makes this challenging, several observations can be made. In the case of the MDA-MB-231 TCs, cells are widely distributed in the extravascular space and often line the abluminal surface of endothelium along and sometimes appear to merge with the endothelium, as observed by overlapping UEA-1 lectin and TC fluorescence signals. TCs were observed directly lining the lumen, closely resembling mosaic vessels or vessel mimicry previously observed *in vivo*^43,44^. Mosaic vessels are hypothesized to form from a loss of CD31 (or UEA1 lectin in this case) reactivity, EC shedding, or rapid vessel proliferation leading to TC entry^43^, and further studies are required to understand their role, if any, in intravasation. Interestingly, a previous *in vitro* TME model using an EC-lined channel often observed mosaic vessels and, in some instances, subsequent vessel collapse^20^, whereas vessel collapse was not observed in the 3D self-assembled vessels in the IntravChip. In our system, we observed a TC in the process of intravasating appearing to squeeze through the endothelium. Whether this occurs at cell-cell junctions or not is not immediately clear from our initial studies. Other questions on the process of intravasation remain, such as what are the primary methods TCs use to enter the circulation, or whether there are specific conditions that promote TC intravasation. While addressing these questions will require additional studies, we have shown here that the IntravChip recapitulates several critical observations previously reported in *in vitro* and *in vivo* studies^20,43,44^.

One of our objectives was to examine the effect of varying types of TMEs (presence of flow, TC density, and TC type) on the vascular morphology and intravasation potential. We observed a significant increase in vessel area coverage in TMEs with flow, while varying TC density or TC type produced little or no effect of the TME vasculature. TC-laden tissues showed a non-significant trend toward higher vessel permeability than TC-free tissues. They also exhibited a notable increase in variability, which could suggest the presence of local, transient regions of higher permeability that may exist where TCs are protruding through the endothelium^2^. Furthermore, we have previously shown that vessel permeability significantly decreases within two days of commencing continuous flow^55^, suggesting that there may be competing effects: a decrease in permeability from continuous perfusion and an increase due to the presence of TCs^16^. Flow significantly increased TC area coverage in the TME, possibly due to improved gas exchange and nutrient delivery. Thus, flow is not only an essential biophysical stimulus, but it also affects the vessel morphology and TC proliferation.

Even in the TC-dense TME in the IntravChip, direct observation of TC intravasation was rare. Prior 3D *in vitro* models relied on either lengthy time-lapse imaging to capture these rare events^18^ or TC migration after transmigrating through the endothelium to identify intravasated TCs^21,17^, motivating the need for a TC collection chamber to directly assess intravasation rates. We observed 100-440 TCs in the collection chamber over five days of flow. This rate, however, may be lower than occurs *in vivo* since our system lacked macrophages, which are known to increase intravasation rates^2,18^. But capturing most of the intravasated TCs in the collection chamber enabled us to observe the significant influence of the presence/absence of flow, TC density in the TME, and TC type on the intravasation rate. We found that flow is essential to collecting the intravasated TCs in the IntravChip. When expressed as a fraction of TCs in the TME, a low TC density produced higher efficiency in less dense tumors. A prior study observed that MDA-MB-231 TCs seeded at a low density (50-70% confluency in 2D) were more invasive across an endothelial monolayer and showed higher intravasation and extravasation rates in zebrafish compared to TCs seeded at a high density (100% confluency in 2D)^56^. The IntravChip captured significant differences in TC invasiveness across TC types, with the highly metastatic lines MDA-MB-231 and MV3 yielding the highest intravasation rates, and the low metastatic line MCF7 yielding the lowest rate. One caveat, however, is that the TC count after 9 days is likely an overestimate of the number of intravasated cells due to proliferation after entering the collection chamber. To address this limitation, future studies could employ single-cell microfluidic impedance cytometry as an alternative method for directly detecting flowing, intravasated TCs^57^.

The IntravChip enables analysis of TC behavior across multiple spatial scales, from cellular morphology to nanoscale chromatin organization. By combining this platform with STORM, we directly visualized heterochromatin architecture in tumor cells undergoing intravasation. Our STORM imaging results show that intravasation drives substantial nanoscale remodeling of heterochromatin in metastatic breast cancer cells. Despite preserved nucleus-wide H3K9me3 abundance, intravasated TCs display smaller, fragmented, and more broadly dispersed heterochromatin nanodomains compared with the compact structures observed in primary TME TCs. This redistribution is consistent with the mechanical stresses associated with TC intravasation, which can deform the nucleus and reorganize chromatin architecture. Such mechanically induced nanoscale chromatin reorganization has been documented in other contexts, including the aberrant chromatin configurations observed under chemo-mechanical perturbations^29,30^, where chromatin structure responded sensitively and dynamically to extracellular mechanical cues. Although our findings contrast with prior studies^47^ reporting increased heterochromatin accumulation following confined migration, these discrepancies likely arise from fundamental differences in experimental context. Whereas earlier models^47^ relied on acute, largely 2D passage through rigid constrictions and short-term analysis, our system captures tumor cells within a prolonged, 3D multicellular microenvironment where they interact with endothelial cells, remodel matrix, and undergo intravasation, more closely recapitulating *in vivo* progression and enriching for a highly migratory subpopulation. Within this physiologically relevant setting, we resolve nanoscale heterochromatin reorganization during intravasation, a critical layer of genome organization that underlies gene regulation^58,59^, highlighting the IntravChip as a powerful platform to study chromatin dynamics in metastasis.

A major utility of preclinical cancer models is their use in drug screening. In this study, we showed that the IntravChip reproduced previously reported *in vivo* and *in vitro* observations of sorafenib’s effect on reducing tumor size^50,51,53^ and intravasation events^16^, validating the IntravChip’s use as a preclinical drug screening platform. Our model also provides several additional insights. First, the 3D vasculature in the IntravChip enables the assessment of anti-cancer drugs, which often have anti-angiogenic functions, on vascular structure and function. Using the IntravChip, a 5 μM dose of sorafenib was sufficient to significantly decrease intravasation events without having adverse effects on the vascular structure, while a 10 μM dose significantly decreased vessel diameter and area coverage, observations which are only possible in models with 3D vasculature. Second, we observed that sorafenib treatment significantly reduced the number of collected intravasated TCs, indicating that it may also reduce TC dissemination, a potentially new mechanism of action. This result corroborates a previous *in vivo* study showing that sorafenib treatment reduced metastases in a rat tumor model^60^. Furthermore, a retrospective study reported a correlation between sorafenib treatment and a decrease in incidence of brain metastases in a subpopulation of patients^61^. Altogether, these results indicate that in addition to reducing the primary tumor size, sorafenib also reduces intravasation. They also demonstrate the potential of the IntravChip as a drug screening platform and the ability to distinguish drug effects on TCs and TME vasculature.

The IntravChip offers several advantages over previous *in vitro* models used to study intravasation, namely this system incorporates 3D vasculature with physiological permeabilities, the system can support a highly dense TC population in the TME while maintaining perfusable vessels, and it enables collection of the intravasated TC population providing a direct measure of intravasation rates. Future iterations of the IntravChip could feature a modified primary TME compartment to accommodate TC spheroids or patient samples, either directly added during the initial seeding phase or introduced later once the vasculature is fully formed. Additional cell types could be included in the primary TME, such as immune cells^62^ or other stromal cells, to recapitulate different types of solid tumors.

## Conclusion

We present a novel microfluidic model of the primary TME that enables collection of intravasated TCs for quantitative downstream analyses. The TME in this system supports a dense TC population with an embedded three-dimensional, perfusable vasculature, recapitulating the close TC-EC interactions observed *in vivo*. Continuous perfusion of the IntravChip enables efficient collection of intravasated TCs and provides physiologically relevant mechanical stimuli. Using this platform, we investigated TMEs with varying TC densities and TC types. Downstream analysis revealed that intravasated TCs exhibit smaller chromatin domain sizes compared to TCs retained within the primary TME. Finally, we validated the IntravChip’s application as a drug screening tool using sorafenib. Together, these results establish the IntravChip as a physiologically relevant platform for mechanistic studies of intravasation and for functional and molecular characterization of metastasis-competent TCs.

## Acknowledgements

We thank the Koch Institute’s Robert A. Swanson (1969) Biotechnology Center for data management support, specifically Charlie Demurjian and Stuart Levine at The Barbara K. Ostrom (1978) Bioinformatics and Computing Facility. We thank Dr. Peter Friedl and Katie Craven for insightful discussions on improving the design of the IntravChip and interpreting the experimental results. MF was supported by an MIT MathWorks Fellowship. EC was supported by an Early Postdoc.Mobility fellowship from the Swiss National Science Foundation (P2EZP2_199914), a postdoctoral fellowship from the Ludwig Center at MIT Koch Institute for Integrative Cancer Research, and a Postdoc.Mobility fellowship from the Swiss National Science Foundation (P500PB_222131). This work was supported by a grant from the National Cancer Institute (U54-CA261694). The funder played no role in study design, data collection, analysis and interpretation of data, or the writing of this manuscript.

