## Supplemental Information for "IntravChip: a vascularized and perfused microfluidic model of the primary tumor microenvironment to collect intravasated tumor cells"

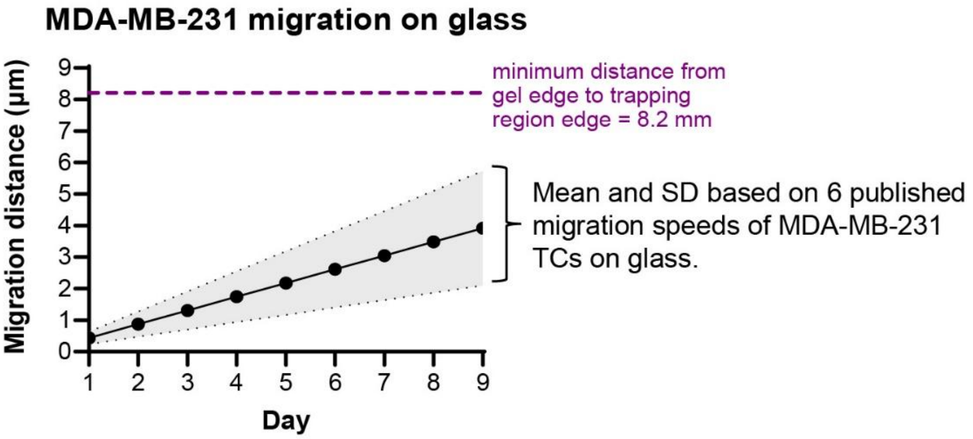


**Supp. Fig. 1. Migration distance of MDA-MB-231 TCs on glass over 9 days.** Migration speeds were taken from published sources[1–5]. The migration distance estimate here assumes that the TCs are migrating along a straight line, indicating the maximum possible distance travelled, although TCs are known to have variability in their direction of migration.

**Supp. Fig. 2. Derivation of the relationship to estimate the minimum radius of the collection chamber.**

The size of the collection chamber was designed such that the residence time of the TCs in the settling chamber was greater than the time required for the TC to settle to the glass based on differences in densities between the TCs and the media.

An estimate of the minimum radius of the collection chamber can be found through a force balance between the drag force on the settling cell,

$$F_d = 3\pi\mu v_s D \quad (1)$$

where $F_d$ is the drag force, $\mu$ is the medium viscosity, $v_s$ is the settling velocity of the TC, and $D$ is the diameter of the TC. Buoyancy acting on a TC suspended in medium is defined as,

$$F_g = \frac{1}{6}\pi D^3 g(\rho_{cell} - \rho_{medium}) \quad (2)$$

where $F_g$ is the buoyancy, $g$ is gravity, $\rho_{cell}$ is the TC density, and $\rho_{medium}$ is the medium density. Setting (1) and (2) equal to one another and rearranging provides a relationship for the settling velocity of the TC,

$$v_s = \frac{1}{18\mu_{medium}} D^2 g(\rho_{cell} - \rho_{medium}) \quad (3),$$

where $\mu_{medium}$ is the viscosity of the medium, $D^2$ is the diameter of the TC, $g$ is gravity, and $\rho_{cell}$ and $\rho_{medium}$ are the densities of the TC and medium, respectively. The characteristic settling time, $t_s$, of the TC is

$$t_s = {}^{h}/_{v_s} \quad (4),$$

where $h$ is the height of the settling chamber. The characteristic time of a TC flowing through a circular collection chamber can be estimated as

$$t_r = {}^{V_{chamber}}/_{Q} \quad (5),$$

where $V_{chamber}$ is the volume of the settling chamber and $Q$ is the volumetric flow rate. For a TC to settle, we require that $t_r > t_s$. Subbing in (4) and (5), we obtain an expression to estimate the minimum radius of the collection chamber,

$$r_{collection\ chamber} > \sqrt{\frac{9\mu Q}{2\pi g r_{cell}{}^2(\rho_{cell} - \rho_{medium})}} \quad (6).$$

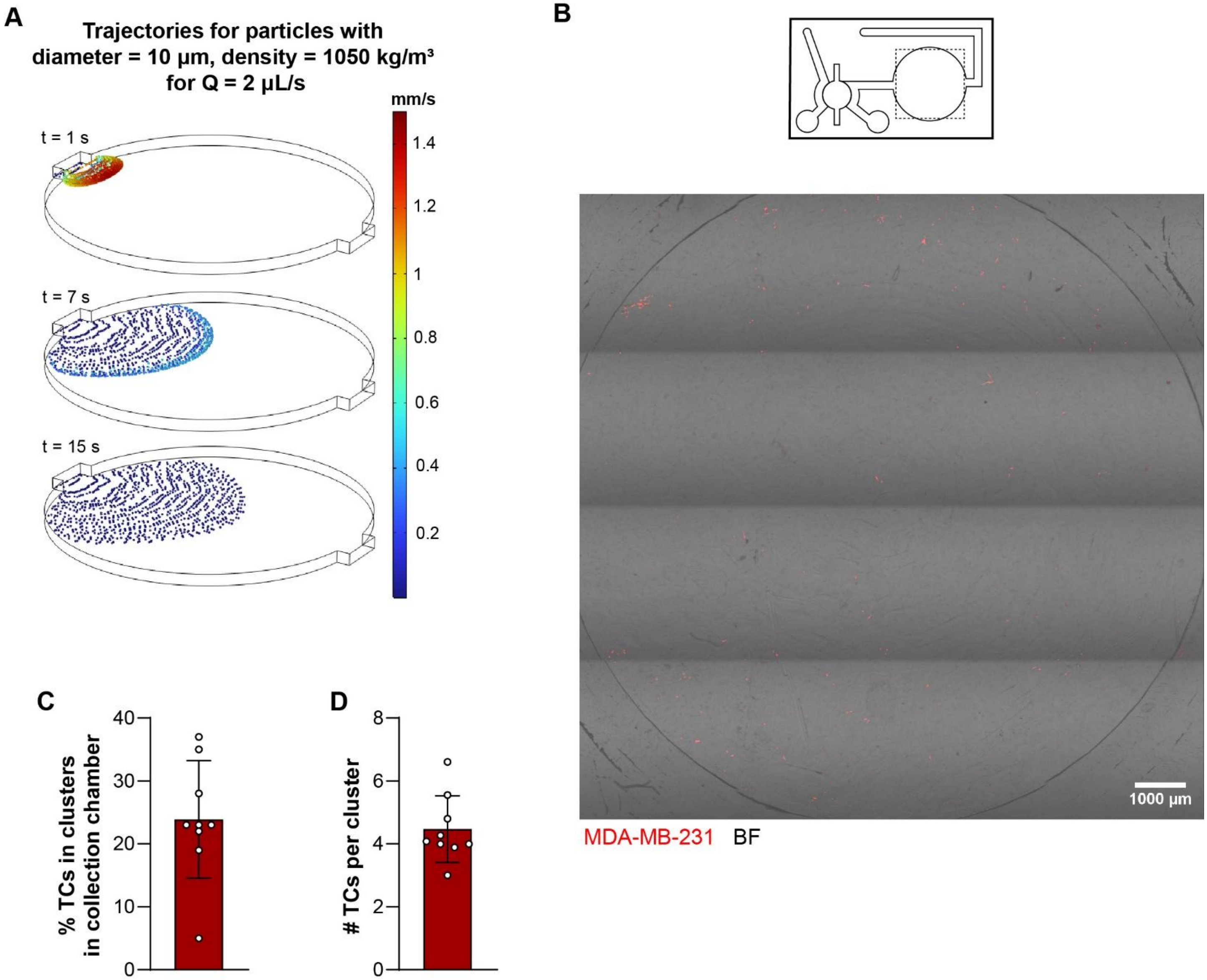


**Supp. Fig. 3. TC distribution in the collection chamber from simulations and experiments at day 9.** (A) Time course of the particle tracing simulation showing the radial distribution of collected particles representing collected TCs. (B) Representative image of the collection chamber with collected TCs. (C) Percentage of TCs in the collection chamber found in clusters of 3 or more TCs. (D) Average number of TCs composing the TC clusters in the collection chamber. Graphs show mean +/- sd. n = 9 devices.

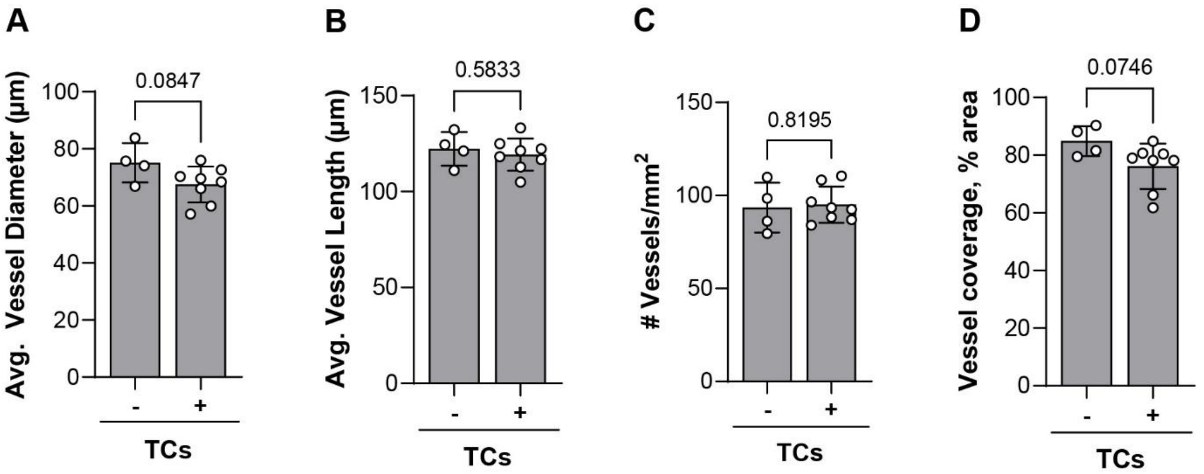


**Supp. Fig. 4. Assessment of vessel morphology in TC-free (-) or TC-laden (+) tissues on day 7.** (A) Average vessel diameter, (B) average vessel length, (C) vessel density, and (D) percent vessel area coverage in the tissues. n = 4-8 devices. Graphs show mean +/- sd.

### Supplementary Information References
